# What to compare and how: comparative transcriptomics for Evo-Devo

**DOI:** 10.1101/011213

**Authors:** Julien Roux, Marta Rosikiewicz, Marc Robinson-Rechavi

## Abstract

Evolutionary developmental biology has grown historically from the capacity to relate patterns of evolution in anatomy to patterns of evolution of expression of specific genes, whether between very distantly related species, or very closely related species or populations. Scaling up such studies by taking advantage of modern transcriptomics brings promising improvements, allowing us to estimate the overall impact and molecular mechanisms of convergence, constraint or innovation in anatomy and development. But it also presents major challenges, including the computational definitions of anatomical homology and of organ function, the criteria for the comparison of developmental stages, the annotation of transcriptomics data to proper anatomical and developmental terms, and the statistical methods to compare transcriptomic data between species to highlight significant conservation or changes. In this article, we review these challenges, and the ongoing efforts to address them, which are emerging from bioinformatics work on ontologies, evolutionary statistics, and data curation, with a focus on their implementation in the context of the development of our database Bgee (http://bgee.org).

## Introduction

Variation in animal or plant morphology is caused by variation in their genomes. Evolutionary developmental biology, or Evo-Devo, aims at understanding the origin of morphological variations, whether between closely related species (e.g., the variation in wings shapes and colors between sister species of butterflies (Brunetti et al., 2001, Beldade et al., 2002, Monteiro et al., 2009)), or between very distant lineages (e.g., the divergence between insect and vertebrate body-plans (Raff, 2000, De Robertis, 2008)), and how genomic variation translates into morphological variation at all scales. Anatomical differences between species are studied in the context of the developmental processes generating species anatomy, and the role of genes in these developmental processes. The involvement of genes and their function in the developmental process is traditionally determined by comparing their expression patterns between species. A particular emphasis is put on genes which exhibit either rapid changes in expression, which have been linked to numerous interesting developmental differences, or deep conservation of expression patterns, which have been show to underlie developmental similarities on unexpectedly large evolutionary scales (Raff, 2000, Carroll, 2005). This has long been done on a small scale, a few genes at a time, using techniques such as *in situ* hybridization or quantitative RT-PCR. With microarrays, then RNA-seq, it has become possible to study expression patterns on a genome-wide scale. Yet application of these techniques to questions from Evo-Devo has lagged somewhat behind other fields. This stands in contrast to the rapid and productive use that was made of the increasingly available genome sequences, illustrated for example by the in-depth study of the evolution of *Hox* genes (Hoegg et al., 2005), or enhancers sequences (Holland et al., 2008).

Although there has been recent progress in the comparison of transcriptomes of closely related species (Gallego Romero et al., 2012, Necsulea et al., 2014) the difficulties in answering many of the questions posed in Evo-Devo remains. How do changes in expression patterns translate to changes in the function of homologous organs? What is the molecular basis for morphological innovation? What can deep conservation of expression tell us about homology relations between distant species? How much of transcriptome evolution is convergent vs. parallel? Here, we discuss some of these issues and emerging solutions. We believe that the next years will be a very fruitful time for the application of transcriptomics to Evo-Devo, and that this will yield important insights into the evolution of development and of anatomy.

## What to compare? The problem of anatomical homology

A major problem in comparing expression patterns on a large scale between species is the definition of what is comparable. In comparative genomics, orthologous genes can be readily identified by sequence comparison, and reliable data are now available from numerous databases (Sonnhammer et al., 2014). But expression patterns are much more complex combinations of genes and of anatomical and temporal patterns. Thus criteria to guide the comparison between these structures are necessary.

The most obvious comparison criterion is homology, as is used for the genes themselves. Homology is most commonly taken in the sense of “historical homology”, a definition in which the common evolutionary origin has to derive from common descent, i.e. the common ancestor of both species possessed the anatomical structure, and both present day species being compared inherited it from this ancestor. But the interpretations of the homology concept have changed with progresses in biology, and different subfields operate with different definitions (Roux et al., 2010). Notably, the “biological homology” concept considers organs homologous if they share a set of developmental constraints (Wagner, 1989). This process-oriented and more mechanistic definition, mostly used in Evo-Devo, encompasses repeated parts in the same organism (e.g., somites), as well as sexually differentiated parts of individuals of the same species (e.g., testis and ovaries), which do not pass the common ancestry criterion, yet are important to our understanding of the relation between development and the evolution of form.

For transcriptome comparisons between very closely related species, such distinctions are usually not an issue: homologous structures are mostly obvious and comply with most definitions of the homology concept. For example, in comparisons among Great Apes, such as between humans and chimpanzees (see (Gallego Romero et al., 2012) for a review), it is mostly trivial to accept that structures with the same name are homologous, and that moreover they essentially perform the same function. But when more distant species are compared, which anatomical structures should be compared becomes less obvious. And it is not always clear whether the most biologically relevant comparisons should involve homologous or functionally equivalent structures. Indeed, some homologies are well established, but correspond to organs which are widely divergent in function and structure (e.g., tetrapod lung and actinopterygian fish swim bladder (Zheng et al., 2011)). For other structures, such as the vomeronasal organ in different tetrapodes, or the ovary between teleost fishes and mammals, similarity of function is clear but homology is debated (for a discussion see (Niknejad et al., 2012)). Similarly, the homology of segmentation mechanisms among bilaterian lineages remains unclear (De Robertis, 2008).

To take into account the complexity of homology on a large scale, we have been developing computational representations of homology, through ontologies (Parmentier et al., 2010, Roux et al., 2010, Niknejad et al., 2012), and using them in the annotation of transcriptome data to anatomical information. The distribution of the ontologies (Noy et al., 2009), and the recent merger of our efforts into Uberon (Mungall et al., 2012, Haendel et al., 2014), allows any comparative transcriptomics or comparative functional genomics project with an interest in homology to built upon this work (e.g., Robinson et al., 2014).

## What to compare? Homologous organs are not always functional equivalents

Other comparisons than homology may be relevant to detect sets of genes whose expression is responsible for the emergence or maintenance of particular functions. It is generally assumed that the expression of core genes responsible for conserved functions should be itself conserved (Levine et al., 2005). But in homologous structures, the signal of functional constraint might be difficult to characterize from the background similarity in gene expression due to their common origin. The task of isolating the common molecular signals responsible for functional similarity could thus be facilitated by comparing non-homologous organs with similarity in function, e.g., tetrapod lung *vs.* fish gills, both used as respiratory organs, or structures used in pregnancies of female mammals *vs*. male Syngnathids (seahorses and pipefishes) (Stölting et al., 2007), or homologous organs that present functions that were not inherited from the common ancestor but have evolved in parallel, e.g., body parts of cichlid fishes which evolved in parallel in African or Central American lakes to perform the same functions in species that colonized similar ecological niches (Elmer et al., 2011).

Many studies have successfully related patterns of phenotypic convergence to patterns of gene expression, at the scale of few genes. Between closely related species, it was notably shown that independent genetic changes could be responsible for regulatory changes, often of homologous genes, leading to the parallel evolution of similar phenotypes (Martin et al., 2013). Examples include the evolution of black spots on wings of different *Drosophila* species, involving the regulation of the *yellow* gene (Gompel et al., 2005, Prud’homme et al., 2006), or the parallel evolution of opsins expression in cichlids (Hofmann et al., 2010, O’Quin et al., 2010). Between marine and freshwater populations of the stickleback fish, differences in the number of armor plates have been mapped to regulatory changes of the *EDA* gene (Jones et al., 2012). Interestingly, convergent patterns of expression changes in the same genes are often caused by nucleotide substitutions affecting the same *cis-*regulatory elements (Elmer et al., 2011).

There are still few transcriptome-wide studies of such convergent anatomical structures, but recent work highlight the diversity of signatures underlying phenotypic convergence. For example the convergent evolution of bacterial photophores of two squid species extends beyond anatomy, to highly convergent transcriptome profiles (Pankey et al., 2014). Whereas the convergent evolution of external genitalia in different amniotes is mirrored in patterns of transcriptomes which relate more to the tissue of origin of these new structures than to the evolution of their functional equivalence (Tschopp et al., 2014). It remains to be seen which of these divergent evolutionary trajectories is more general, and which factors could lead to deep reorganization of transcriptomes with the evolution of new functions.

At larger evolutionary distances, such as between insects and vertebrates, there are few unambiguous homologous anatomical structures between species, and most shared functions are the result of convergence or parallel evolution. But fascinating, and unexpected, examples of gene expression patterns similarity were uncovered by Evo-Devo studies. The most famous example is probably the case of vertebrate and arthropods eyes, which are not homologous, but express similar transcription factors during their developmental cascade (including *Pax6* (Halder et al., 1995)). The concept of “deep homology” (Shubin et al., 2009) has been proposed to describe the relation between structures which share some homology in their development, notably in their key patterning genes, share also similarity of function and of structure, yet are not homologous at the anatomical level, certainly not in the sense of historical homology (deep homology is formalized in our HOM ontology (Roux et al., 2010) as a special case of parallelism, which is a case of homoplasy). Other examples illustrate such a co-option of common, reusable toolkits, such as the developmental program of arthropod and vertebrate appendages, implicating the transcription factor *Dll*/*Dlx* (Nielsen et al., 2003). Interestingly, this gene is also expressed in the distal tips of “outgrowths” in diverse animals, for example the horns of beetles (Moczek, 2006).

Conversely, convergent or parallel phenotypes can evolve with non-homologous gene expression. For example, the convergent evolution of coat color in different subspecies of beach mice probably evolved through changes in expression of different genes (Manceau et al., 2010). More striking, homologous structures are not always patterned by conserved expression patterns. For example the vulva of different species of nematodes are patterned by non-homologous pathways (Schlager et al., 2006) and the proteins expressed in the crystalline lens of different vertebrate species are unrelated (Piatigorsky et al., 1991).

Overall, the correspondence between homology at the levels of gene expression and of anatomy is complex (Hall, 1994), yet with proper experimental design can be very informative. Genes can be co-opted to conduct the same function in non-homologous structures, while vastly divergent expression can underlie homologous structures. Determining which anatomical comparisons are likely to be informative in the comparison of transcriptomes between species is not easy, especially between very distant species, and requires good anatomical and developmental knowledge. A challenge of future Evo-Devo transcriptomics is to integrate such knowledge with genome-scale studies to provide relevant insight into the relation between the evolution of organ function and homology.

## What to compare? The problem of heterochrony

For developmental processes, as for anatomy, we need to define what is comparable if we are to perform meaningful transcriptome comparisons. But unlike for anatomical structures, there are no clear homology relations between developmental stages. This is mostly because of heterochrony, differences in the relative timing of developing organs between species. For example, the human and rat developmental stages at which heart reaches the same developmental milestone might differ, and other structures in these two species can reach their developmental milestones in a different order (detailed in Jeffery et al., 2002). Thus, a pair of developmental stages between two species will rarely present similarly advanced development for all the organs or tissues forming the embryos.

Accordingly, clear examples of heterochrony were found for the expression of several gene pathways in a whole body comparison of closely related Xenopus species (Yanai et al., 2011). In a more anatomically targeted study focusing on the development of human, chimpanzee and rhesus macaque prefrontal cortex, as many as 71% of genes were found to change in timing of expression (Somel et al., 2009). In both of these examples, closely related species were compared. For more distant species, very broad developmental periods can be compared, to diminish the impact of heterochrony and ensure some level of comparability. For example, in Comte et al. (2010), we calculated the conservation of expression of orthologous genes between zebrafish and mouse for seven broad developmental stages, which was sufficient to detect significant differences between early development (up to neurula) and late development, but lacked fine resolution. These broad bilaterian “meta-stages” are available in our database of gene expression evolution Bgee (Bastian et al., 2008). Alternatively, all-against-all developmental stages can be compared (Kalinka et al., 2010, Parikh et al., 2010, Irie et al., 2011). This approach has allowed to identify interesting patterns of expression conservation or divergence over ontogeny, notably a minimum temporal divergence between Drosophila species at mid development (Kalinka et al., 2010, Gerstein et al., 2014, Li et al., 2014).

## Can we compare levels of expression between species on a large-scale?

In classical small-scale Evo-Devo studies gene expression patterns have been typically compared in terms of presence or absence (Carroll, 2005). Yet expression levels can be key in understanding changes in ontogeny. For example, differences in metamorphosis among vertebrates, including heterochronic shifts, seem to be driven in large part by changes in the expression level of key genes (Laudet, 2011). Similarly, a comparison of expression levels in mammalian organs identified numerous modules of genes with conserved tissue-specific expression, but whose absolute expression levels shifted in particular taxonomic groups, e.g., primates (Brawand et al., 2011, Necsulea et al., 2014). More broadly the comparison of transcriptomes between Xenopus species found that variation in expression levels constituted most transcriptome changes (Yanai et al., 2011). It is thus relevant to consider expression levels as a primary evolving phenotype.

Microarrays allowed the first large-scale studies of the evolution of gene expression between species, but their analysis in an evolutionary context poses specific problems. Most microarray datasets whose comparison would be interesting were generated by different studies carried out separately in different species. Absolute expression levels are thus difficult to compare directly in such cases because they suffer from important batch effects. There can also be large biases in signal between microarray platforms and technologies used in different species, for example because the hybridization strength depends on probes and on gene sequence composition. Thus, to detect conserved or divergent levels of expression between species, post-processing approaches are often used. It is for example possible to compare lists of co-expressed genes, or modules, derived individually in each species (Bergmann et al., 2003, Stuart et al., 2003, Lu et al., 2009, Piasecka et al., 2012a). It is also possible to compare lists of differentially expressed genes between different conditions in each species, or to compare lists of functional categories of genes enriched for differentially expressed genes in each species (Lu et al., 2009).

Correlative methods have been used to compare the hybridization signals across species. The most commonly used methods are Pearson or Spearman correlations (gene expression is considered conserved if the orthologs are strongly correlated across different conditions or tissues between species) and Euclidian distance (Su et al., 2004, Liao et al., 2006, Xing et al., 2007, Parikh et al., 2010). However, the results of these approaches have been shown to differ greatly when different measures or data normalization schemes are used, and they behave differently for tissue-specific genes or broadly expressed (housekeeping) genes (Piasecka et al., 2012b), which complicates their interpretation. Additionally, correlative methods rarely take into account the variability of expression levels within species.

Direct comparing of hybridization signal is more legitimate when technical problems of microarrays in comparative studies are minimized or circumvented, for example, when the hybridization of samples from different species is performed within the same study, on the same microarray platform (Ranz et al., 2003, Rifkin et al., 2003, Nuzhdin et al., 2004, Ometto et al., 2011). Unfortunately, such an approach only works for the comparison of very closely related species where the hybridization of cDNAs on probes of the microarray is not perturbed by the low nucleotide divergence in the sequence of orthologous genes (Lu et al., 2009). Otherwise, probes that do not match exactly all orthologous target genes have to be ignored (Khaitovich et al., 2005, Kalinka et al., 2010). Customized multi-species microarrays have also been designed to eliminate the sequence mismatch effects. Samples from two species are competitively hybridized to the probes and intensity ratios are estimated by averaging signal on probes from the different species (Gilad et al., 2005, Vallee et al., 2006, Oshlack et al., 2007). However these approaches are dependent on having a high quality genome annotation for each species in order to locate sequence differences and to account for their potential effect.

RNA-seq is now allowing major progresses in describing gene expression variation between species. This technique has a larger dynamic range than microarrays, and can also be used to study differences in exon usage and alternative splicing (Gallego Romero et al., 2012). Importantly, it allows the study of non-model species in the absence of a sequenced genome (Grabherr et al., 2011, Perry et al., 2012), or when the genome sequence is of poor quality. The advantages of RNA-seq allow more straightforward direct comparisons of expression levels between species, and interesting insights have been provided by the first evolutionary studies using this technology (Blekhman et al., 2010, Brawand et al., 2011, Perry et al., 2012, Khan et al., 2013).

When expression levels are directly compared across species, heuristic approaches have mostly been used to identify patterns of conservation or of directional selection (Fraser, 2011). Expression divergence between species is compared to the divergence within biological replicates of the same species. This procedure is similar to the McDonald and Kreitman test (MK test) of positive selection, widely used to characterize selection on protein-coding sequences (McDonald et al., 1991). Genes under stabilizing selection display limited expression variation both within and between species, whereas genes under directional selection in a given lineage display a shift in mean expression between species, while maintaining a low variation within species. Of note, alternative explanations for the latter pattern are possible, such as a relaxation of selective constraints (although this scenario should be accompanied with an increase of within species variation). Progress has recently been made towards the development of formal models to test for positive selection on expression levels (Rohlfs et al., 2014). Still, these approaches cannot control for differences in environmental input to gene regulation between species (Gallego Romero et al., 2012).

Even though many technological challenges have been overcome in the past years, the comparison of expression levels between species holds numerous remaining difficulties (Dunn et al., 2013), particularly when the species compared are distant. First, expression levels measured in a given tissue are a mix of expression levels of the cell types constituting this tissue. Tissues of different species often display differences in cellular composition, notably in the proportion and localization of different cell types. This factor can result in artificial differences in expression levels and bias the comparison between species (Pantalacci et al., in press). For example, brain, blood or pancreatic samples were shown to typically vary substantially in their cellular composition between even closely related species (Hill et al., 2005, Magalhaes et al., 2010, Steiner et al., 2010).

Second, a basic sources of measurement noise both in microarray and RNA-seq transcriptomic studies comes from alternative splicing (Barbosa-Morais et al., 2012, Merkin et al., 2012, McManus et al., 2014). Alternative-splicing patterns have been shown to be evolve fast and in consequence are mostly species-specific. A difference of the isoforms expressed between species can affect the measure of expression level because of differences in length (e.g., longer transcripts produce more reads in RNA-seq) and in exon structure (e.g., microarray probes may be designed to bind only some exons). Although RNA-seq facilitates the reconstruction of the pool of all isoforms for each gene, which then could ideally be examined in isolation, this task still proves to be very challenging (Hayer et al., 2014).

Third, the heterogeneity of annotation may bias expression comparison between species. Gene models for orthologs may contain fragments that are not orthologous. A solution to this problem is to carefully align the nucleotide sequences of orthologs sets to compare, and restrict further analysis only to fragments that are strictly orthologous (Blekhman et al., 2010, Brawand et al., 2011). The difference in the genomic context of orthologous genes between species can also be an issue. Different genomes contain different numbers of paralogous genes and pseudogenes, which both may be sources of spurious reads or unspecific hybridization background. This also challenges the comparison of genomic regions whose sequences are unique in the genome of some species but not others, leading to differential mappability of RNA-seq reads.

Although several recent papers illustrate the power of comparative RNA-seq and other functional genomics approaches between close species, and even try to extend them to the comparison of distant species (Boyle et al., 2014, Chen et al., 2014, Gerstein et al., 2014, Li et al., 2014), these studies remain limited in the anatomical and developmental complexity which is covered. In consequence their applications to fundamental Evo-Devo questions remains limited. A notable study focused on the evolution of gene expression in distant Dictyostelium species, and was facilitated by the conserved morphology of these species (Parikh et al., 2010). But to link functional genomics to more fine questions of Evo-Devo, more detailed transcriptome profiles are needed, which will in turn raise the issues of homology and heterochrony which have so far been circumvented.

## Data sets for comparative transcriptomics are increasingly available and integrated

Transcriptomics data, whether from microarrays or RNA-seq experiments, is increasing exponentially in public databases (Rustici et al., 2013). Although it is mostly generated to answer biomedical and other practical questions, it covers an increasing number of organs, tissues, developmental stages, and species, of interest for Evo-Devo studies. The challenge is to identify, recover and organize the relevant data among the tens of thousands of tumor and cell line samples. While there are several ongoing efforts to organize transcriptome data (notably the Gene Expression Atlas (Kapushesky et al., 2010) or the Gene Expression Barcode (McCall et al., 2014); see also (Rung et al., 2013)), we will focus here on our database Bgee (http://bgee.org) (Bastian et al., 2008), which is the only effort to our knowledge which focuses on Evo-Devo concerns: inter-species comparisons using anatomical homology, detailed developmental stage annotation, and present/absent calls for different expression data types. We provide an overview of some major transcriptomic datasets of interest for Evo-Devo, whether they are already integrated into Bgee or will be in the near future.

Bgee was the first database to integrate together RNA-seq, microarray and *in situ* hybridization data. Bgee release 13 (December 2014) will integrate more than 500 Illumina RNA-seq libraries from 14 animal species: human (*H. sapiens*), chimpanzee (*P. troglodytes*), gorilla (*G. gorilla*), macaque (*M. mulatta*), mouse (*M. musculus*), rat (*R. norvegicus*), pig (*S. scrofa*), cow (*B. taurus*), opossum (*M. domestica*), platypus (*O. anatinus*), chicken (*G. gallus*), lizard (*A. carolinensis*), frog (*X. tropicalis*) and worm (*C. elegans*). A high proportion of these RNA-seq samples comes from a few large-scale comparative studies, such as Brawand et al. (2011), which contains samples from brain, cerebellum, heart, kidney, liver and testis from nine species of placental mammals (great apes, rhesus macaque and mouse), marsupials (grey short-tailed opossum) and monotremes (platypus); Barbosa-Morais et al. (2012), which complemented the above dataset with samples from 5 tissues of lizard and frog; and Merkin et al. (2012), which generated RNA-seq samples from 9 tissues from 4 mammals (rat, mouse, cow and macaque), and one bird (chicken). The value of such comparative RNA-seq datasets is reflected by their rapid reuse in independent comparative studies (Barbosa-Morais et al., 2012, Gokcumen et al., 2013, Reyes et al., 2013, Hong et al., 2014, Washietl et al., 2014, Yang et al., 2014). For model organisms, these RNA-seq are completed by microarray data; the high number of collected experiments, and available methods for processing and analysis, still make microarrays a valuable resource for meta-analyses. Bgee release 13 will integrate more than 12,000 Affymetrix arrays, mapped on more than 400 anatomical structures in five model species: mouse, human, zebrafish (*D. rerio*), fruit fly (*D. melanogaster*) and worm.

All microarray and RNA-seq datasets come from public repositories such GEO (Gene Expression Omnibus)(Barrett et al., 2013) or ArrayExpress (Rustici et al., 2013) and are carefully annotated by Bgee curators to appropriate anatomical structures and developmental stages classes of the ontologies (Haendel et al., 2014). All selected datasets and samples were collected in normal (non-treated) conditions, and from wild-type animals. This latter condition is critical for evolutionary studies, where expression should be as much as possible relevant to the evolutionary history and wild-type selective pressures experienced by the species compared.

It is clear that the integration of datasets for non-model species obtained with RNA-seq technique is much easier than equivalent microarray datasets prepared using different technologies and often even with in-house designed platforms. Such data, from diverse species, is increasingly available, and much of it is not yet integrated into Bgee. For example, a newly emerging source of transcriptomic data from many primate species is the non-human primate reference transcriptome resource (NHPRTR). Their full RNA-seq dataset consists of 157 libraries across 14 species or subspecies: Chimpanzee (*P. troglodytes*), Indochinese Cynomolgus macaque (*M. fascicularis*), Mauritian Cynomolgus Macaque (*M. fascicularis*), Gorilla (*G. gorilla*), Japanese macaque (*M. fuscata*), Chinese Rhesus macaque (*M. mulatta*), Indian Rhesus macaque (*M. mulatta*), Pig-tailed macaque (*M. leonina*), Olive baboon (*P. anubis*), Common marmoset (*C. jacchus*), Ring-tailed lemur (*L. catta*), Sooty mangabey (*C. atys*), Squirrel monkey (*Saimiri sp.*), Mouse lemur (*Microcebus sp.*), and up to 14 different tissues (Pipes et al., 2013).

Additionally, the ENCODE and modENCODE consortia have recently released large amounts of RNA-seq data from mouse, human, worm and fly sampling numerous developmental stages, tissue and cell types, much of them highly relevant to Evo-Devo studies. Notably, more than 140 RNA-seq worm samples and 250 fly samples cover their development at great resolution (35 developmental stages for worm and 30 developmental stages for fly) (Graveley et al., 2011, Gerstein et al., 2014, Li et al., 2014). Although the majority of the human transcriptomic data from the ENCODE project comes from cancer or immortalized cell lines, which limits its use for evolutionary analysis, more than 200 human RNA-seq libraries were recently used along with worm and fly data for the identification of evolutionary conserved transcription modules between these distantly related species (Gerstein et al., 2014).

On a final note, while RNA-seq and microarrays provide genome-wide information, they often lack the fine resolution of *in-situ* hybridizations, which are mostly small-scale. Yet, there exist a few large-scale projects aimed at generating *in-situ* hybridizations for every gene of a species at different developmental time points, for example for mouse (Diez-Roux et al., 2011) or for zebrafish (Thisse et al., 2004). Apart from these systematic efforts, a collection of many small-scale studies can sum up to a large valuable resource, as reflected by the success of the Gene Ontology functional annotations (du Plessis et al., 2011). The abundance of *in situ* hybridizations integrated from published studies now provides an almost genomic overview of expression patterns in some species (e.g., Kassahn et al., 2009). Bgee integrates *in situ* hybridization data from 4 databases: BDGP for fruit fly (Hammonds et al., 2013), MGI for mouse (Smith et al., 2014), Xenbase for frog (Bowes et al., 2010) and ZFIN for zebrafish (Bradford et al., 2011). In the current release of Bgee there are 23754 genes with expression information extracted from 231864 *in situ* hybridization images, produced in 35072 independent experiments. Unlike for microarrays and RNA-seq data, the primary extraction and annotation of information from papers is done by the curators of each of these respective databases, while Bgee provides mapping to the bilaterian anatomy ontology Uberon (Haendel et al., 2014), and to ontologies of developmental stages. These data cover anatomical complexity in an extremely detailed manner.

Thus overall there exist abundant transcriptome and gene expression data, which are relevant to the questions posed by Evo-Devo, and modern database efforts make these data increasingly available for genome-wide investigation into evolutionary developmental biology.

## Future directions

A challenge as well as an opportunity of comparative transcriptomics for Evo-Devo is relating large scale quantitative trends with morphological observations. The main limitation of comparative approaches so far is that, while it is relatively easy to establish significant conservation of expression, it is very difficult at a large scale to characterize significant changes of evolutionary relevance. Expression can differ because of many experimental reasons, from environmental conditions to measurement errors, as well as because of low stabilizing selection. Yet it can also differ because of developmental innovations, convergent evolution, or other evolutionary scenarios of interest to understanding morphological variety. In the absence of a good baseline, similar to the synonymous substitution rate of protein-coding gene codon models (Yang, 2006), calling such relevant changes in transcriptomic studies remains a major challenge. In the end, the evolutionary scenarios that are the most difficult to unveil are often the most fascinating, which motivates further methodological work into evolutionary transcriptomics.

Another important challenge is establishing causality: mechanistic causality, as in determining which gene expression changes determine changes in morphology; and evolutionary causality, as in determining which expression patterns are fixed for their role in morphological innovations or constraints. For this, we will probably need to obtain more transcriptomic data of variation both within and between species, and combine it with other functional genomics information, such as transcription factor binding patterns (ChIP-seq) and large-scale mutant screens.

Despite the challenges and limitations, we feel that we are at a privileged moment in the long pursuit of the goal of linking genome evolution to the evolution of form, with its constraints and its adaptive roles. The progress of RNA-seq is unlocking the function of the genomes of many species beyond the classical model organisms, new transcriptomics techniques are arriving which promise to combine genomic scale with anatomical precision (Battich et al., 2013, Lee et al., 2014), and the improvement of ontologies and bioinformatics methods allow us to increasingly take advantage of these genomic datasets to answer long standing questions of Evo-Devo.

## Acknowledgements

J.R. is supported by Marie Curie fellowship PIOF-GA-2010-273290. Work on Bgee and comparative transcriptomics in the M.R.-R. lab is supported by the Swiss Institute of Bioinformatics, by the Swiss National Science Foundation [grants number 31003A 133011/1 and 31003A_153341/1], by SystemsX.ch project AgingX, and by Etat de Vaud.

